# Inflammation in Alzheimer’s disease: do sex and APOE matter?

**DOI:** 10.1101/741777

**Authors:** Paula Duarte-Guterman, Arianne Y. Albert, Amy M. Inkster, Cindy K. Barha, Liisa A.M. Galea, on behalf of the Alzheimer’s Disease Neuroimaging Initiative

## Abstract

Alzheimer’s disease (AD) disproportionately affects females with steeper cognitive decline and more neuropathology compared to males, which is exacerbated in females carrying the APOEε4 allele. The risk of developing AD is also higher in female APOEε4 carriers in earlier age groups (aged 65-75), and the progression from cognitively normal to mild cognitive impairment (MCI) and to AD may be influenced by sex. Inflammation is observed in AD and is related to aging, stress, and neuroplasticity, and although studies are scarce, sex differences are noted in inflammation. The objective of this study was to investigate underlying physiological inflammatory mechanisms that may help explain why there are sex differences in AD and APOEε4 carriers. We investigated, using the ADNI database, the effect of sex and APOE genotype (non-carriers or carriers of 1 and 2 APOEε4 alleles) and sex and diagnosis (cognitively normal (CN), MCI, AD) on CSF (N= 279) and plasma (N= 527) markers of stress and inflammation. We found CSF IL-16 and IL-8 levels were significantly lower in female non-carriers of APOEε4 alleles compared to males, whereas levels were similar between the sexes among carriers of APOEε4 alleles. Furthermore, females had on average higher levels of plasma CRP and ICAM1 but lower levels of CSF ICAM1, IL-8, IL-16, and IgA than males. Carrying APOEε4 alleles and diagnosis (MCI and AD) decreased plasma CRP in both sexes. Sex differences in inflammatory biomarkers support that the underlying physiological changes during aging differ by sex and tissue origin.

## INTRODUCTION

Alzheimer’s disease (AD) is a neurodegenerative disease characterized by severe cognitive decline [1]. Risk factors for AD include modifiable risk factors such as sociocultural or lifestyle factors (e.g., education, marital status, exercise), chronic stress exposure [2], and medical conditions (diabetes, obesity, and cardiovascular disease) [3–5]. Non-modifiable lifetime risk factors for AD include age, female sex, and APOE genotype [6]. However, research on the effects of biological sex on risk for AD is equivocal and may depend on geographic location (reviewed in [4,7,8]). Nevertheless, females with AD show greater cognitive decline [9–11] and neuropathology compared to males (faster brain atrophy rates, neurofibrillary tangles; [10,12–15]). Intriguingly, the presence of APOEε4 alleles increases the risk to develop AD in females compared to males at an earlier age (aged 65-75; [16]), and accelerates neuropathology and cognitive decline more so in females than in males [10,11,14,17–19], indicating that the APOE genotype interacts with sex on various factors related to AD. However, there is limited research into the role of sex and its interaction with APOE genotype in the possible mechanisms underlying AD. Understanding why females in general and female APOEε4 carriers have a higher burden of the disease is important for the development of tailored treatments. Biomarkers are highly sought after to predict disease onset and progression and to understand the underlying mechanisms of diseases in order to develop or improve treatments.

Chronic low grade inflammation is a hallmark of AD, as evidenced by increased expression of proinflammatory cytokines in the brains of AD patients (not analyzed by sex), which can exacerbate AD pathology [20–22]. There is, however, increasing evidence that there are sex differences in immune responses in healthy adults with females mounting a stronger response compared to males after an acute challenge [23,24]. In response to an endotoxin, females have higher levels of pro-inflammatory plasma cytokines (TNF-α and IL-6) while males have higher plasma levels of anti-inflammatory IL-10 [23,25]. In addition, aging affects the immune system differently in males and females, with females having higher genomic activity for adaptive cells and males having activity for monocytes and inflammation [26]. Although limited, there is evidence that sex differences in systemic inflammation are associated with greater AD pathology [27] but not cognitive decline in normal aging [28]. Specifically, higher C-reactive protein (CRP) levels in blood beginning in midlife are associated with higher brain amyloid levels later in life in healthy males, but not in healthy females [27]. To our knowledge, very few studies have stratified by sex and APOE genotype or sex and diagnosis of cognitive status on potential biomarkers of AD, including inflammation.

Sex differences in inflammatory biomarker systems may also differentially affect neuroplasticity [29,30], which is reduced in AD and correlates with cognitive decline [31,32]. In addition, peripheral cortisol, the main stress hormone in humans, is elevated in AD [33] and is associated with higher amyloid levels in the brain [34], a reduction in hippocampal volume, and cognitive impairment in older individuals [35] that may depend on MCI status [36]. Peripheral cortisol is also associated with elevated pro-inflammatory cytokines [23,37]. However, it is not known how sex differences in markers of inflammation (e.g., cytokines, immunoglobulins, CRP, intercellular adhesion molecule, ICAM1), and stress hormones (cortisol) may be related to sex differences in AD.

Using the ADNI database, we conducted exploratory analyses examining sex differences in CSF and plasma physiological biomarkers, inflammation and stress related, and how these may be affected by APOE genotype (non-carriers or carriers of APOEε4 alleles), and dementia status (cognitively healthy (CN), MCI, AD). We tested the hypothesis that females have higher levels of inflammation and stress hormones compared to males and these levels are disproportionately affected by the presence of APOEε4 alleles and AD diagnosis.

## METHODS

### ADNI database

Data used in the preparation of this article were obtained from the Alzheimer’s Disease Neuroimaging Initiative (ADNI) database (adni.loni.usc.edu). The ADNI was launched in 2003 as a public-private partnership, led by Principal Investigator Michael W. Weiner, MD. The primary goal of ADNI has been to test whether serial magnetic resonance imaging (MRI), positron emission tomography (PET), other biological markers, and clinical and neuropsychological assessment can be combined to measure the progression of mild cognitive impairment (MCI) and early Alzheimer’s disease (AD). For up-to-date information, see www.adni-info.org. Data used in this article were downloaded on or before Jan 16, 2019. Inclusion and exclusion criteria of participants [38,39] were the same for the two datasets analysed in the current study (biomarkers in CSF and plasma), and general procedures are detailed online (http://adni.loni.usc.edu/methods/documents/). Briefly, cognitively normal (CN) participants had normal memory function based on education-adjusted scores on the Wechsler Memory Scale Logical Memory II and a Clinical Dementia Rating (CDR) of 0. Amnestic late MCI (LMCI) participants had objective memory loss (measured by education-adjusted scores from Wechsler Memory Scale Logical Memory II), a CDR of 0.5, preserved daily activities, and absence of dementia. All AD participants met NINCDS/ADRDA Alzheimer’s Criteria and a CDR of 0.5 or 1.0.

To address our research questions, we used two separate datasets from the ADNI database: CSF biomarkers and plasma biomarkers (Table 1). Although the datasets do not overlap completely, within the plasma-CSF datasets there is an overlap of 85% (i.e., 85% of individuals with CSF biomarker data also had plasma levels of biomarkers). This is an exploratory study of these variables on sex by APOE genotype and sex by diagnosis and we discuss the limitation of these overlapping datasets below.

**Table 1.** Demographic and clinical information for all ADNI participants subdivided by sex. Participants with measured biomarkers in (A) cerebrospinal fluid (CSF) and (B) plasma. We collapsed APOE genotype into two groups: (1) participants carrying any ε4 alleles (homozygous ε4/ε4 and heterozygous ε4/-) and (2) participants with no ε4 risk alleles (-/-). In the two subdata sets, females were significantly younger and had fewer years of education than males. In data set A (but not B), there was a trend for the proportion of female and male participants in each of the diagnosis to be different (P=0.051) with more females (27.5 % compared to 21.8%) diagnosed with AD, more females cognitively normal (32.1% compared to 22.9%), and fewer females diagnosed with LMCI compared to males (40.4% compared to 55.3%). The proportion of female and male participants carriers and non-carriers of APOEε4 alleles was not significantly different in any of the two datasets analysed. 85% of individuals with CSF biomarkers (A) had also plasma biomakers (B). CN, cognitively normal; LMCI, late mild cognitive impairment; AD, Alzheimer’s disease.

| A. CSF |  |  |  |  | B. Plasma |  |  |  |
| --- | --- | --- | --- | --- | --- | --- | --- | --- |
|  |  | Sex |  |  |  | Sex |  |  |
|  | Total | Female | Male | P-value | Total | Female | Male | P-value |
|  | No. 279 | No. 109 | No. 170 |  | No. 527 | No. 196 | No. 330 |  |
| Age |  |  |  |  |  |  |  |  |
| Mean (SD) | 75.15 (±6.86) | 73.75 (±6.69) | 76.04 (±6.83) | 0.007 | 74.75 (±7.40) | 73.79 (±7.63) | 75.32 (±7.21) | 0.051 |
| Education (years) |  |  |  |  |  |  |  |  |
| Mean (SD) | 15.69 (±2.95) | 14.68 (±2.74) | 16.34 (±2.90) | < 0.0001 | 15.57 (±3.04) | 14.94 (±2.89) | 15.95 (±3.07) | < 0.0001 |
| Ethnicity |  |  |  |  |  |  |  |  |
| White | 267 (95.70%) | 103 (94.50%) | 164 (96.47%) | 0.55 | 498 (94.68%) | 186 (94.90%) | 312 (94.55%) | 0.27 |
| Not White<sup>†</sup> | 12 (4.30%) | 6 (5.50%) | 6 (3.53%) |  | 28 (5.32%) | 10 (5.10%) | 18 (5.45%) |  |
| Baseline diagnosis |  |  |  |  |  |  |  |  |
| CN | 74 (26.5%) | 35 (32.1%) | 39 (22.9%) | 0.051 | 40 (7.6%) | 19 (9.7%) | 21 (6.4%) | 0.16 |
| LMCI | 138 (49.5%) | 44 (40.4%) | 94 (55.3%) |  | 378 (71.9%) | 132 (67.3%) | 246 (74.5%) |  |
| AD | 67 (24.0%) | 30 (27.5%) | 37 (21.8%) |  | 108 (20.5%) | 45 (23.0%) | 63 (19.1%) |  |
| APOEε4 allele number |  |  |  |  |  |  |  |  |
| 0 | 134 (48.03%) | 51 (46.79%) | 83 (48.82%) | 0.81 | 243 (46.20%) | 90 (45.92%) | 153 (46.36%) | 0.93 |
| 1 or 2 | 145 (51.97%) | 58 (53.21%) | 87 (51.18%) |  | 283 (53.80%) | 106 (54.08%) | 177 (53.64%) |  |
| Cortisol (ng/mL) |  |  |  |  |  |  |  |  |
| Mean (SD) | 16.05 (±6.04) | 14.92 (±6.01) | 16.78 (±5.96) | 0.008 | 2.17 (±0.13) | 2.16 (±0.13) | 2.17 (±0.13) | 0.16 |
| C reactive protein (ug/mL) |  |  |  |  |  |  |  |  |
| Mean (SD) | -2.83 (±0.56) | -2.77 (±0.64) | -2.87 (±0.51) | 0.23 | 0.12 (±0.54) | 0.21 (±0.55) | 0.07 (±0.52) | 0.003 |
| CD40 antigen (ng/mL) |  |  |  |  |  |  |  |  |
| Mean (SD) | -0.65 (±0.12) | -0.66 (±0.10) | -0.64 (±0.14) | 0.12 | -0.12 (±0.13) | -0.12 (±0.13) | -0.12 (±0.14) | 0.87 |
| Interleukin 16 (pg/mL) |  |  |  |  |  |  |  |  |
| Mean (SD) | 0.91 (±0.18) | 0.87 (±0.17) | 0.94 (±0.19) | 0.004 | 2.55 (±0.15) | 2.54 (±0.15) | 2.55 (±0.16) | 0.34 |
| Interleukin 3 (ng/mL) |  |  |  |  |  |  |  |  |
| Mean (SD) | -2.22 (±0.32) | -2.28 (±0.29) | -2.17 (±0.34) | 0.001 | -1.65 (±0.29) | -1.65 (±0.29) | -1.65 (±0.30) | 0.97 |
| Interleukin 6 receptor (ng/mL) |  |  |  |  |  |  |  |  |
| Mean (SD) | -0.01 (±0.15) | -0.02 (±0.14) | -0.00 (±0.15) | 0.30 | 1.46 (±0.14) | 1.48 (±0.14) | .45 (±0.13) | 0.02 |
| Interleukin 8 (pg/mL) |  |  |  |  |  |  |  |  |
| Mean (SD) | 1.68 (±0.15) | 1.64 (±0.11) | 1.70 (±0.16) | 0.001 | 1.02 (±0.19) | 1.02 (±0.21) | 1.01 (±0.18) | 0.1 |
| Intercellular adhesion molecule 1 (ng/mL) |  |  |  |  |  |  |  |  |
| Mean (SD) | 0.96 (±0.44) | 0.83 (±0.33) | 1.04 (±0.48) | 0.0001 | 2.01 (±0.15) | 2.04 (±0.14) | 2.00 (±0.15) | 0.03 |
| Immunoglobulin A (mg/mL) |  |  |  |  |  |  |  |  |
| Mean (SD) | -2.54 (±0.31) | -2.68 (±0.26) | -2.45 (±0.31) | < 0.0001 | 0.61 (±0.23) | 0.60 (±0.23) | 0.62 (±0.22) | 0.21 |

P-values are from Wilcoxon rank sum tests for continuous variables and Fisher's exact tests for categorical variables. <sup>†</sup>Includes self-reported Black, Asian, American Indian/Alaskan, and >1 ethnicity.

### Statistical Methods: Inflammatory markers

We included all ADNI participants that had inflammatory markers measured in CSF (N = 279) and plasma (N = 527) listed in Tables 1. Data included in our analyses were: demographics (age, years of education, and ethnicity), baseline diagnosis (cognitively normal, CN; late MCI, LMCI; or AD), and number of APOEε4 alleles. We collapsed APOE genotype into two groups: (1) participants carrying any ε4 alleles (homozygous ε4/ε4 and heterozygous ε4/-) and (2) participants with no ε4 risk alleles (-/-). Plasma and CSF samples from the ADNI study were collected in CN, LMCI, and AD participants at baseline in the morning after an overnight fast. Processing, aliquoting and storage were performed according to the ADNI Biomarker Core Laboratory Standard Operating Procedures. Inflammatory markers were measured using a commercially available multiplex proteomic panel (Human Discovery Multi-Analyte Profile; Luminex xMAP) developed by Rules-Based Medicine (Austin, TX), that measures a variety of markers including cytokines, metabolic markers, and growth factors. We initially chose biomarkers available in plasma involved in inflammation and immune responses (cytokines, immunoglobulins, CRP, and ICAM1) and stress (cortisol; Table 2). We analysed the same biomarkers in CSF (however IgE and IL-18 are not available in CSF). The protocols used to quantify plasma and CSF analytes are described in Craig-Schapiro et al. [40] and Hu et al. [41]. We used the ADNI quality-controlled data for plasma and CSF provided by the ADNI Consortium. For plasma IL-16, we removed one outlier that was more than two times lower than the 25^th^ percentile in the plasma data. Sensitivity analysis with the outlier present suggested that it was disproportionately influencing the results.

**Table 2.**
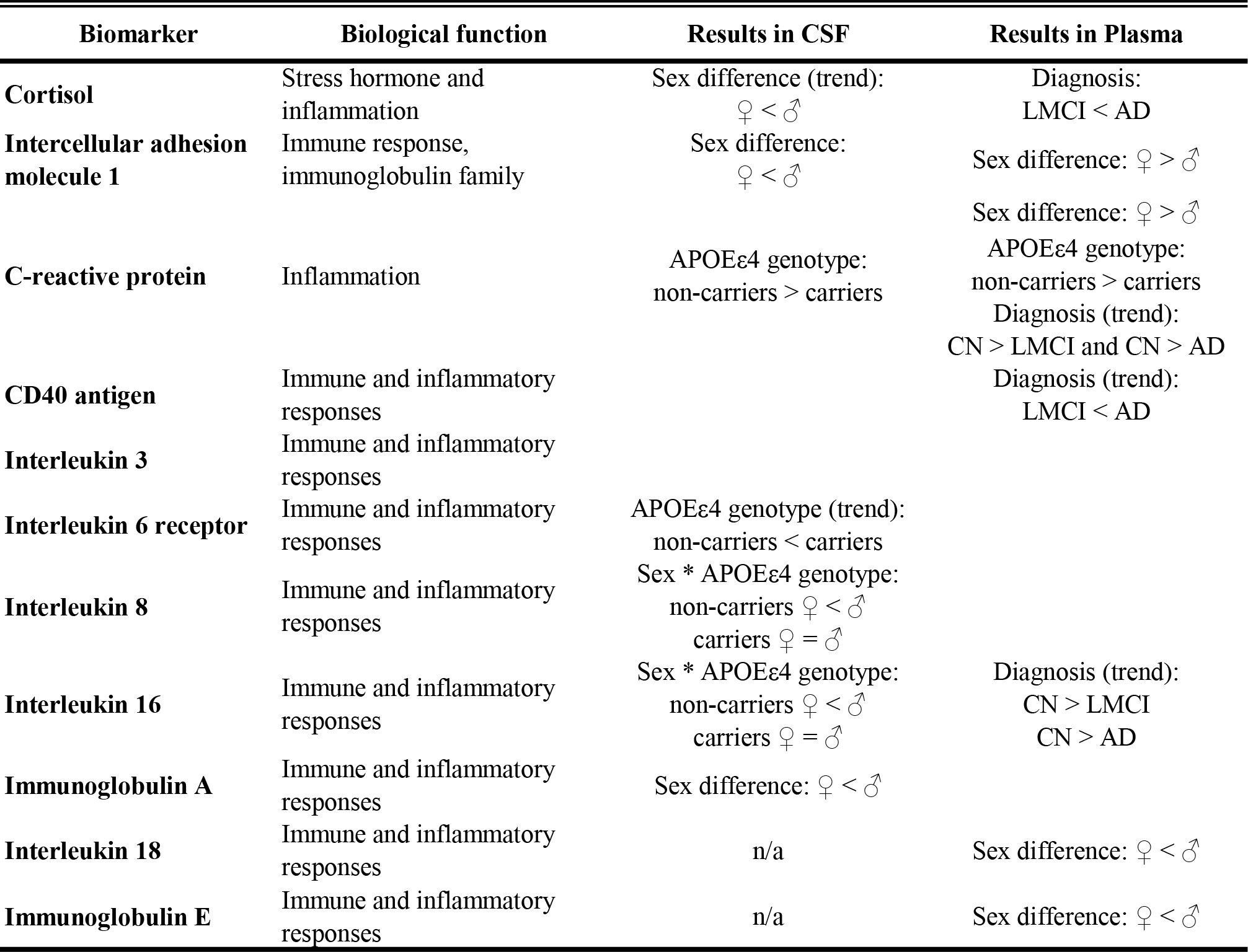
List of biomarkers analysed in the current study with their main biological function and main finding in the CSF and plasma. Main effects of sex (sex difference), APOEε4 genotype (non-carriers or carriers), and diagnosis (CN, cognitively normal; LMCI, late mild cognitive impairment; AD, Alzheimer’s disease) and interaction between sex and APOEε4 genotype (sex * APOEε4 genotype) are shown. Significant effects are adjusted P≤0.05 and trends are adjusted P≤0.08. See results for details. n/a - not available

We compared all available data for each study variable between the sexes using the Wilcoxon rank sum test for continuous variables and Fisher’s exact test for categorical variables. Nonparametric tests are standard for comparing variables where the distribution is unknown or expected to be non-normal. We used general linear models to determine the relationships between (1) sex and APOE genotype (non-carriers or carriers of APOEε4 alleles) or (2) sex and baseline diagnosis as predictor variables, and biomarkers as dependent variables. Due to the limited sample size, we were not able to study sex, APOE genotype, and baseline diagnosis in one model. All models included age and education as covariates. Initially, all models included an interaction between sex and presence of APOEε4 alleles or sex and baseline diagnosis; if this interaction was not significant, it was removed from the model to estimate the main effects of sex and APOE genotype or diagnosis. Significance was based on the likelihood ratio test, and all P-values for comparisons of sex and either APOE genotype or diagnosis for all outcomes combined were corrected for multiple testing using the Benjamini-Hochberg false discovery rate method with the family-wise error rate set to 0.05 [42]. In total, three P-values per dependent variable were included in each set of models (interaction term and main effects of sex and APOE or diagnosis) resulting in 27 P-values corrected in CSF (9 dependent variables) and 33 P-values corrected in plasma (11 dependent variables; Supplementary Tables S1 to S4) for each of the two models (sex and APOE and sex and diagnosis). Significant interaction terms were followed up using pairwise simple-effects tests with Benjamini-Hochberg P-value correction. A subset of participants with CSF measurements had corresponding plasma measurements (N= 237 total, N= 88 females and N=149 males). For each biomarker, we calculated Pearson’s correlation coefficients between CSF and plasma levels in males and females separately. We then compared these correlations using the Fisher r-to-Z transformation and Z-test using the method by Zou [43]. We report significance differences (adjusted P≤0.05) and trends (adjusted P≤0.08). All regression analyses were carried out in R v3.5.1 [44].

## RESULTS

### Demographic information

Table 1 gives a summary of the variables for the participants with: CSF biomarkers (Table 1A; N=279), plasma biomarkers (Table 1B; N=527). Given the differences in sample sizes, we performed demographic analyses on the two datasets. Females were younger than males in the CSF (P<0.01) and plasma data set (P=0.051). In the two datasets, females had fewer years of education than males (Ps<0.0001). Thus, we used age and education level as covariates in the analyses. Although there were no sex differences in distribution of APOEε4 alleles in any of the two datasets (all Ps>0.4), the proportion of participants in each of the diagnosis categories was marginally different for females and males in the CSF dataset (P=0.051; Table 1 A) but not in the plasma dataset (P>0.1; Table 1 B).

### Sex and presence of APOEε4 alleles were associated with changes in inflammatory markers

Our first aim was to investigate whether sex and APOE genotype interact to influence inflammation using biomarkers, which we analysed separately in CSF and plasma (Supplementary Table S1 and S3, respectively). Caution should be noted as inflammatory signalling can differ depending on tissue examined [45,46].

For inflammatory markers measured in CSF, only IL-16 and IL-8 elicited a significant interaction between sex and APOE genotype (P= 0.016 and P=0.035, respectively; Table 3). CSF IL-16 and IL-8 levels were significantly lower in females non-carriers of APOEε4 alleles compared to males (both P’s<0.001), whereas levels were similar between the sexes in carriers of APOEε4 alleles (P’s>0.9; Fig 1 A and B). Furthermore, in females with APOEε4 alleles, IL-16 was significantly higher than in non-APOEε4 female carriers (P=0.050), while a trend was observed in males (P=0.062). Whereas for IL-8, males with APOEε4 alleles had lower levels of IL-8 compared to males with no APOEε4 alleles (P=0.014) but there was no difference in females (P>0.3). Regardless of sex, CSF CRP levels were lower in carriers of APOEε4 alleles compared to non-carriers (main effect of genotype: P=0.009; Table 3; Fig 1 C). There was a trend for an increase in IL-6 receptor levels in APOEε4 carriers regardless of sex compared to non APOEε4 carriers (main effect of genotype: P=0.071; Table 3). Lastly females had significantly lower CSF levels of IgA and ICAM1 and a trend for lower CSF cortisol levels compared to males (main effect of sex: P<0.001; P=0.009, and P=0.070, respectively; Table 3). There were no other significant main or interaction effects on any other CSF biomarkers.

**Figure 1.**
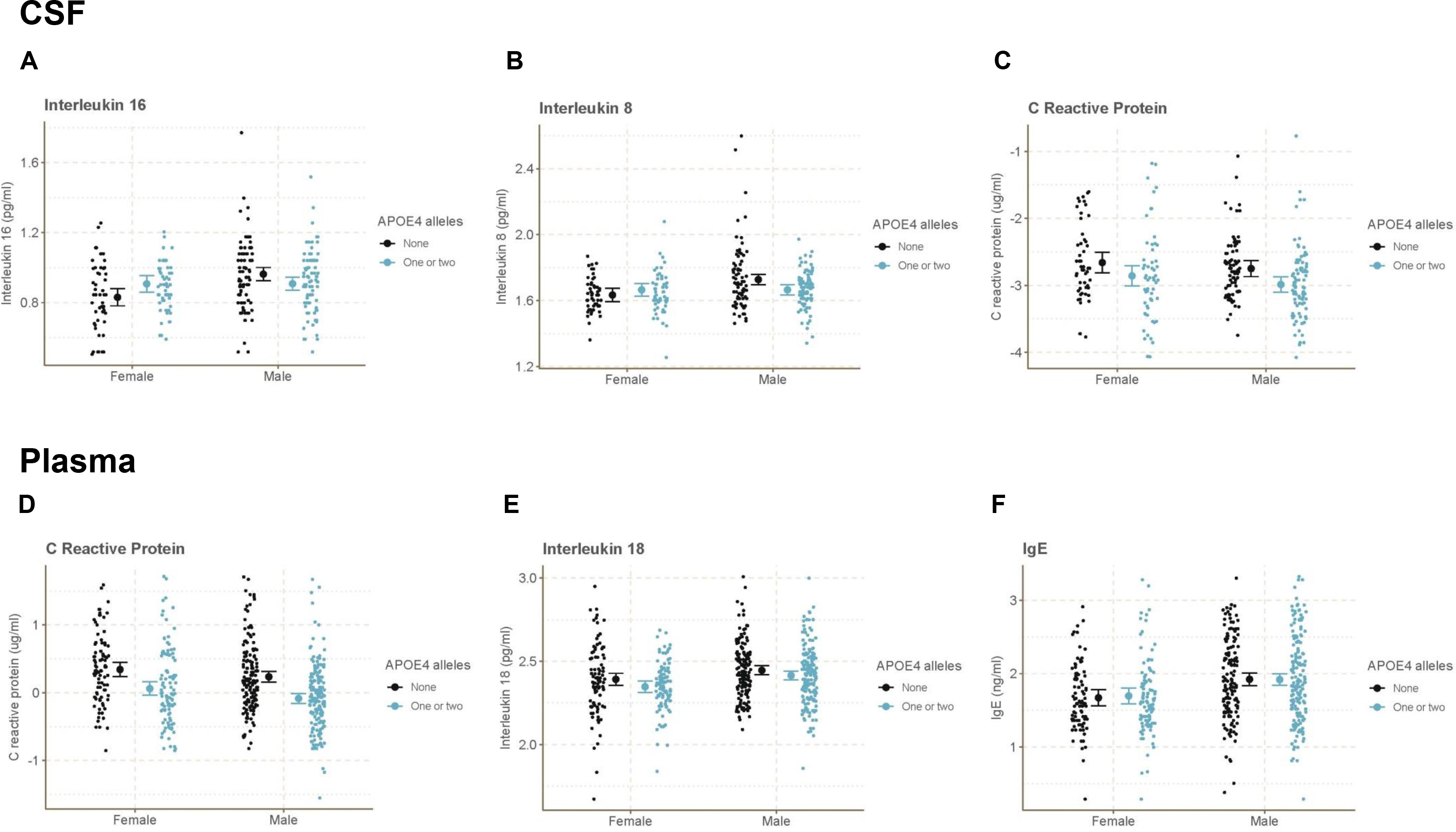
Marginal mean (±95% confidence interval) of **CSF levels** of A. IL-16 (pg/ml), B. IL-8 (pg/ml), C. C-reactive protein (CRP; μg/ml), and **plasma levels** of D. CRP (CRP; μg/ml), E. IL-18 (pg/ml), and F. IgE (ng/ml) in ADNI participants by sex and presence or absence of APOEε4 alleles (none or 1 and 2 alleles).

**Table 3.** Linear regression results for models with sex and APOE genotype (non-carriers or carriers of 1 or 2 APOEε4 alleles). P-values are for overall tests and are FDR-adjusted. Only shown are the models with significant associations (adjusted P ≤0.05) and trends (adjusted P≤0.08). All model summaries are available in Supplementary Table S1.

| Predictors | Cortisol Cortisol ng/ml |  |  | C Reactive Protein ug/ml |  |  | Interleukin 16 pg/ml |  |  | Interleukin 6 receptor ng/ml |  |  | Interleukin 8 pg/ml |  |  | Immunoglobulin A mg/ml |  |  | Intercellular Adhesion Molecule ng/ml |  |  |
| --- | --- | --- | --- | --- | --- | --- | --- | --- | --- | --- | --- | --- | --- | --- | --- | --- | --- | --- | --- | --- | --- |
|  | Estimates (CI) | p | adjusted p | Estimates (CI) | p | adjusted p | Estimates (CI) | p | adjusted p | Estimates (CI) | p | adjusted p | Estimates (CI) | p | adjusted p | Estimates (CI) | p | adjusted p | Estimates (CI) | p | adjusted p |
| (Intercept) | 1.43 (-7.00 – 9.87) |  |  | -3.03 (-3.84 – -2.23) |  |  | 0.42 (0.15 – 0.68) |  |  | -0.27 (-0.48 – -0.06) |  |  | 1.33 (1.11 – 1.54) |  |  | -2.91 (-3.35 – -2.48) |  |  | -0.23 (-0.83 – 0.38) |  |  |
| AGE (years) | 0.22 (0.12 – 0.32) |  |  | 0.01 (-0.00 – 0.02) |  |  | 0.01 (0.00 – 0.01) |  |  | 0.00 (0.00 – 0.01) |  |  | 0.00 (0.00 – 0.01) |  |  | 0.00 (-0.00 – 0.01) |  |  | 0.01 (0.01 – 0.02) |  |  |
| EDUCATION (years) | -0.21 (-0.45 – 0.03) |  |  | -0.01 (-0.03 – 0.02) |  |  | -0.01 (-0.01 – 0.00) |  |  | -0.00 (-0.01 – 0.00) |  |  | 0.00 (-0.00 – 0.01) |  |  | 0.00 (-0.01 – 0.01) |  |  | -0.00 (-0.02 – 0.02) |  |  |
| Male (ref = Female) | 1.72 (0.25 – 3.19) | 0.022 | **0.07** | -0.11 (-0.25 – 0.03) | 0.126 | 0.22 | 0.13 (0.07 – 0.19) |  |  | 0.02 (-0.02 – 0.05) | 0.372 | 0.5 | 0.09 (0.04 – 0.14) | <0.001 |  | 0.21 (0.14 – 0.29) | <0.001 | **<0.001** | 0.18 (0.07 – 0.28) | 0.001 | **0.009** |
| APOE4 1 or 2 alleles (ref = 0 alleles) | 0.82 (-0.55 – 2.19) | 0.241 | 0.35 | -0.22 (-0.35 – -0.09) | 0.001 | **0.009** | 0.08 (0.01 – 0.14) |  |  | 0.04 (0.00 – 0.07) | 0.025 | **0.071** | 0.03 (-0.02 – 0.09) | 0.261 |  | -0.00 (-0.07 – 0.07) | 0.898 | 0.97 | 0.08 (-0.02 – 0.18) | 0.128 | 0.22 |
| Interaction: Male by 1 or 2 alleles |  |  |  |  |  |  | -0.13 (-0.22 – -0.05) | 0.003 | **0.016** |  |  |  | -0.09 (-0.16 – -0.02) | **0.008** | **0.035** |  |  |  |  |  |  |
| Observations | 279 |  |  | 279 |  |  | 279 |  |  | 279 |  |  | 279 |  |  | 279 |  |  | 279 |  |  |
| R<sup>2</sup> / adjusted R<sup>2</sup> | 0.095 / 0.082 |  |  | 0.055 / 0.042 |  |  | 0.122 / 0.106 |  |  | 0.046 / 0.032 |  |  | 0.092 / 0.076 |  |  | 0.123 / 0.111 |  |  | 0.105 / 0.092 |  |  |

For biomarkers measured in plasma, there were no significant interactions between sex and APOE genotype (Supplementary Table S3). However, females had higher plasma CRP levels (main effect of sex: P=0.048; Fig 1 D) and ICAM1 (trend for a main effect of sex: P=0.051) compared to males and significantly lower levels of IL-18 (main effect of sex: P=0.001; Fig 1 E) and immunoglobulin E (IgE: main effect of sex: P<0.001; Fig 1 F) compared to males. Furthermore, plasma CRP decreased in carriers of APOEε4 alleles compared to non-carriers (main effect of genotype: P<0.001; Fig 1 D).

### Sex and baseline diagnosis were associated with changes in inflammatory markers

We next tested whether sex and baseline diagnosis status (CN, LMCI, and AD) influenced CSF and plasma biomarkers of inflammation (Supplementary Table S2 and S4, respectively). There were no significant interactions between sex and diagnosis for any of the tested variables in CSF (Table 4 and Supplementary Table S2) or plasma (Supplementary Table S4). For CSF levels, females had significantly lower levels of IgA (main effect of sex: P<0.001) and ICAM1 (main effect of sex: P=0.026) and a trend for lower IL-16 levels (main effect of sex: P=0.055) compared to males (Table 4; Fig 2 A-C), but we did not observe any significant main effects of diagnosis for any CSF variable.

**Table 4.** Linear regression results for models with sex and baseline diagnosis (CN, cognitively normal; LMCI, late mild cognitive impairment; AD, Alzheimer’s disease). Only shown are the models with significant associations (adjusted P≤0.05) and trends (adjusted P≤0.08). P-values are for overall tests and are FDR-adjusted. All model summaries are available in Supplementary Table S2. There were no significant interactions between diagnosis and sex.

| *Predictors* | **Interleukin 16 pg/ml** |  |  | **Immunoglobulin A mg/ml** |  |  | **Intercellular Adhesion Molecule ng/ml** |  |  |
| --- | --- | --- | --- | --- | --- | --- | --- | --- | --- |
|  | *Estimates (CI)* | *p* | *adjusted p* | *Estimates (CI)* | *p* | *adjusted p* | *Estimates (CI)* | *p* | *adjusted p* |
| (Intercept) | 0.51 (0.26 – 0.77) |  |  | -2.92 (-3.35 – -2.48) |  |  | -0.19 (-0.80 – 0.42) |  |  |
| AGE (years) | 0.01 (0.00 – 0.01) |  |  | 0.00 (-0.00 – 0.01) |  |  | 0.01 (0.01 – 0.02) |  |  |
| EDUCATION (years) | -0.01 (-0.01 – 0.00) |  |  | 0.00 (-0.01 – 0.01) |  |  | -0.00 (-0.02 – 0.02) |  |  |
| Male (ref = Female) | 0.06 (0.02 – 0.11) | 0.007 | **0.055** | 0.21 (0.14 – 0.29) | <0.001 | **<0.001** | 0.17 (0.06 – 0.28) | 0.002 | **0.026** |
| Diagnosis (ref = CN) |  | 0.4 | 0.61 |  | 0.99 | 0.99 |  | 0.54 | 0.67 |
| LMCI | 0.01 (-0.05 – 0.06) |  |  | 0.00 (-0.08 – 0.09) |  |  | 0.06 (-0.06 – 0.18) |  |  |
| AD | -0.03 (-0.09 – 0.03) |  |  | -0.01 (-0.11 – 0.09) |  |  | 0.02 (-0.12 – 0.16) |  |  |
| Observations | 279 |  |  | 279 |  |  | 279 |  |  |
| R<sup>2</sup> / adjusted R<sup>2</sup> | 0.099 / 0.082 |  |  | 0.124 / 0.107 |  |  | 0.101 / 0.085 |  |  |

**Figure 2.**
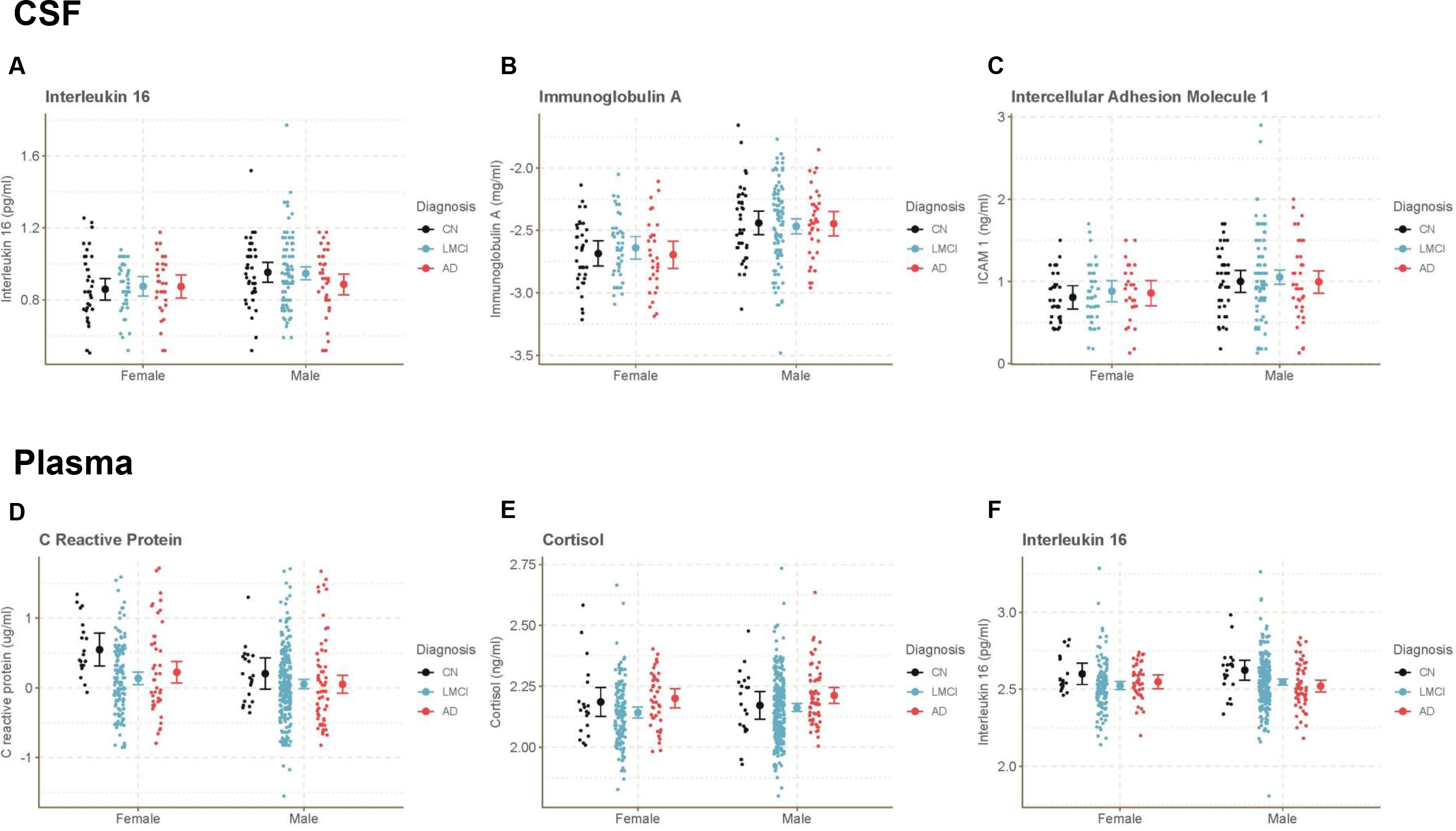
Marginal mean (± 95% confidence interval) of **CSF levels** of A. IL-16 (pg/ml), B. IgA (mg/ml), C. Intercellular adhesion molecule (ICAM1; ng/ml), and **plasma levels** of D. C-reactive protein (CRP; μg/ml), E. cortisol (ng/ml), and F. IL-16 in ADNI participants by sex and diagnosis (CN, cognitively normal; LMCI, late mild cognitive impairment; and AD, Alzheimer’s disease).

In plasma, we found that females had lower levels of IgE (main effect of sex: P<0.001) and IL-18 compared to males (main effect of sex: P=0.004) and trends for females to have higher levels of ICAM1 (main effect of sex: P=0.056) and CRP (main effect of sex: P=0.056; Fig. 2 D) compared to males. In addition, we found diagnosis significantly influenced plasma cortisol (main effect of baseline diagnosis: P=0.01) with lower levels in LMCI compared to AD (P<0.001; Fig 2 E). We found trends for diagnosis to influence plasma IL-16, CRP, and CD 40 levels (main effect of diagnosis: P=0.054, P=0.056; P=0.067). Plasma IL-16 (P’s=0.006) and CRP (P=0.006 and P=0.02) levels were lower in LMCI and AD compared to CN (Fig 2 D and F). For plasma CD 40, levels were lower in LMCI compared to AD (P=0.01; Supplementary Table S4). In summary, although we detected associations between sex and diagnosis and various biomarkers, we did not find evidence of a sex and diagnosis interaction on any variables examined.

### Correlations between cerebrospinal and plasma levels of biomarkers were mostly positive

The results for inflammatory markers in plasma did not always match results in CSF (Supplementary Tables S2 and S4). We therefore investigated the relationship between plasma and CSF biomarkers in males and females (Table 5, Fig 3). Perhaps surprisingly, we found the majority of biomarkers were significantly positively correlated between plasma and CSF levels in both males and females. These significant positive correlations included CRP (males, r=0.793; females r=0.860; P’s<0.0001), IL-6 receptor (males, r=0.459; females r=0.493, P’s<0.0001), IgA (males, r=0.705; females r=0.529; P’s<0.0001), and cortisol in both sexes (males, r=0.176; females, r=0.327; P=0.032 and 0.002, respectively). IL-16 was significantly correlated in females (r=0.290, P=0.006) but only a trend in males (r=0.156, P=0.058). Plasma and CSF levels of ICAM1 and CD 40 were positively correlated in males only (r=0.231, P=0.005 and r=0.374, P<0.0001, respectively) whereas plasma and CSF IL-3 levels were negatively correlated in females only (r=-0.246, P=0.021; Fig 3). There were significant sex differences, favoring males, in the strength of correlation between the sexes for CD 40 (P=0.01) and IgA (P=0.03), with trends for sex differences, favouring males in ICAM1 (P=0.06) and favouring females in IL-3 (P=0.06).

**Table 5.** Pearson’s correlations between plasma and CSF levels of the biomarkers analysed in the current study separetly in males and females. Differences in the correlations were determined using confidence intervals. Significant correlations and differences between correlations are P≤0.05 and trends are P≤0.08

|  | Correlation (r) in<br>Males n = 149 | P-value | Correlation (r) in<br>Females n = 88 | P-value | Difference (95% CI) | P-value |
| --- | --- | --- | --- | --- | --- | --- |
| Cortisol | 0.176 | **0.032** | 0.327 | **0.002** | -0.151 (-0.388-0.101) | 0.24 |
| C reactive protein | 0.793 | **< 0.0001** | 0.860 | **< 0.0001** | -0.067 (-0.149-0.019) | 0.12 |
| CD40 antigen | 0.374 | **< 0.0001** | 0.016 | 0.88 | 0.358 (0.103-0.606) | **0.01** |
| IL-16 | 0.156 | **0.058** | 0.290 | **0.006** | -0.134 (-0.376-0.121) | 0.30 |
| IL-3 | 0.001 | 0.989 | -0.246 | **0.021** | 0.247 (-0.016-0.493) | **0.06** |
| IL-6 receptor | 0.459 | **< 0.0001** | 0.493 | **< 0.0001** | -0.034 (-0.232-0.179) | 0.75 |
| IL-8 | 0.138 | 0.093 | 0.287 | **0.007** | -0.149 (-0.392-0.107) | 0.25 |
| IgA | 0.705 | **< 0.0001** | 0.529 | **< 0.0001** | 0.176 (0.012-0.361) | **0.03** |
| ICAM1 | 0.231 | **0.005** | -0.021 | 0.849 | 0.252 (-0.011-0.507) | **0.06** |

**Figure 3.**
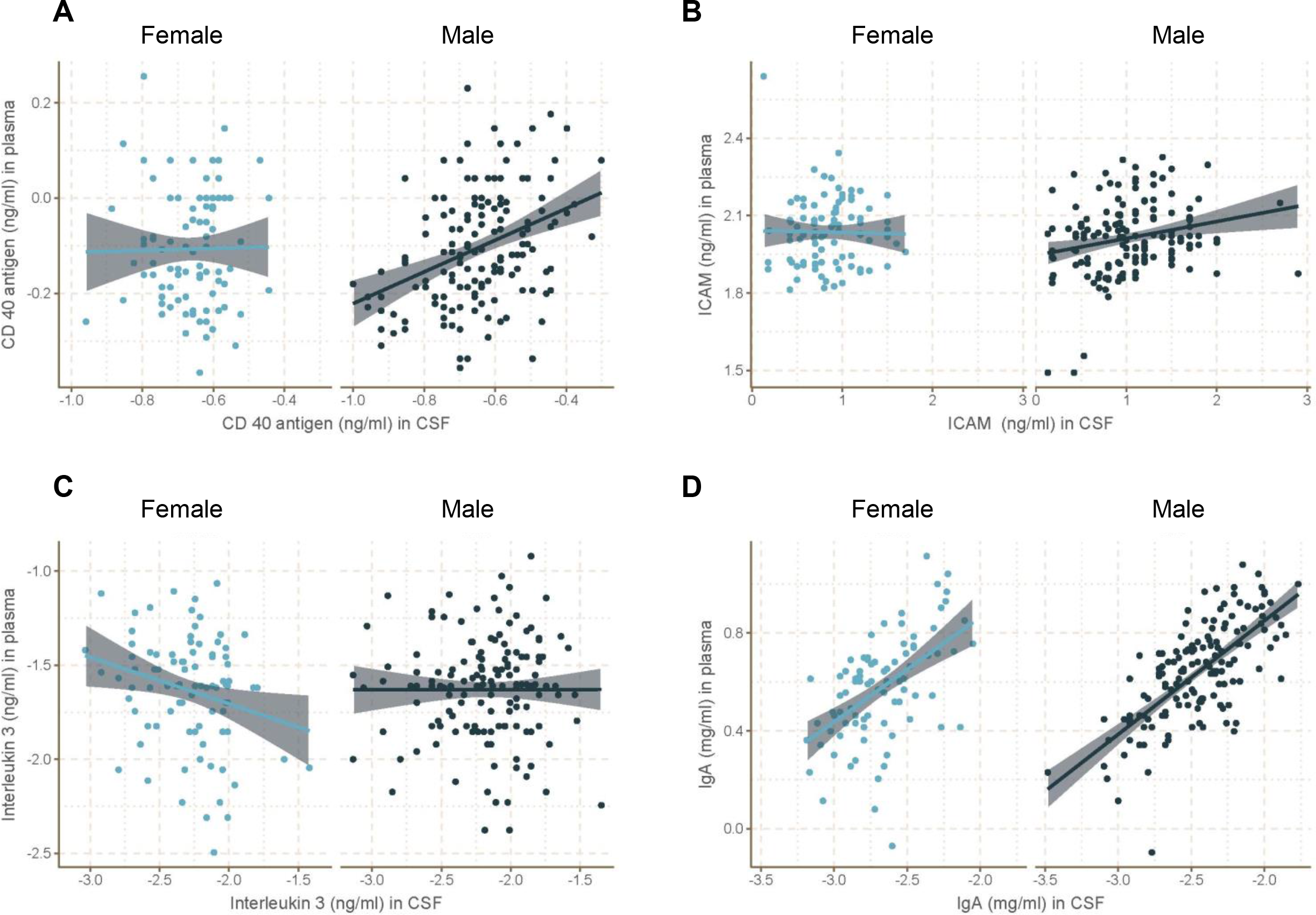
Correlations between plasma and CSF levels of A. CD 40, B. ICAM1, C. IL-3, and D. IgA in males and females separately. CD 40 and ICAM1 were positively correlated in males while IL-3 was negatively correlated in females. IgA was more strongly correlated in males compared to females (see Table 5 for details).

## DISCUSSION

In the present study using ADNI data from CN, LMCI, and AD participants we found interactions between sex and APOE genotype (but not between sex and diagnosis) on CSF and plasma levels of IL-8 and IL-16 (see Table 2 for summary of the results). CSF levels of IL-8 and IL-16 were on average lower in female APOEε4 non-carriers compared to males but similar levels were found between the sexes in APOEε4 allele carriers. Regardless of sex, the APOEε4 allele was associated with decreased levels of CSF and plasma CRP. Sex differences were seen in inflammatory markers, regardless of diagnosis or genotype, as females had lower CSF cytokines (IL-16, IL-18), CSF ICAM1, CSF and plasma immunoglobulins (IgA, IgE), and plasma IL-18. However, tissue (CSF, plasma) mattered for results for certain inflammatory markers (ICAM1 and to a lesser extent CRP) as females had higher plasma CRP and ICAM1 compared to males, opposite to what was found in CSF. Despite these differences in outcomes between plasma and CSF biomarker analyses, plasma and CSF levels were positively correlated for cortisol, CRP, IL-6 receptor, IgA in both sexes, whereas IL-16, and IL-8 were correlated in females and CD 40 and ICAM1 were correlated in males, indicating good consistency between CSF and plasma levels of these biomarkers. Intriguingly, IL-3 stood out from all these biomarkers with a negative correlation between CSF and plasma levels in females only. Males exhibited significantly stronger correlations between plasma and CSF levels for CD 40 and IgA compared to females. Sex and APOE genotype differences in CSF and plasma inflammatory markers suggest differences in underlying physiology that may affect aging and the progression of AD and this should be considered in future studies. Researchers should be cautioned to use sex as a biological variable in all analyses.

### Sex interacted with presence of APOEε4 alleles to affect levels of IL-8 and IL-16

In this study, we found that sex interacted with APOE genotype to influence CSF IL-16 and IL-8. CSF IL-16 and IL-8 levels were lower in females with no APOEε4 alleles compared to males, but no sex differences in these cytokine levels were detected in participants carrying APOEε4 alleles. Our results suggest that presence of APOEε4 alleles can modulate CSF (and potentially plasma) cytokine levels in a sex-dependent way. The APOE protein can regulate transcription in vitro [47] and APOE4, but not APOE3, increases levels of IL-6 and IL-8 in vitro [48]. In the current study, we found that the sex differences in IL-16 and IL-8 levels disappeared in carriers of APOEε4 alleles. One possibility is that the APOE4 protein regulates cytokine levels differently in males and females. IL-16 has been implicated in AD [49] and plasma IL-16 levels decreased with diagnosis (in males and females; current study) and AD severity (analysis without regard to sex; [50]). On the other hand, levels of IL-8 were not affected by diagnosis, consistent with a meta-analysis of cytokines in AD [22]. It is unclear what the impact of regulation of CSF cytokine levels by sex and APOEε4 has on AD symptoms or pathology, however given that females with APOEε4 alleles are disproportionally affected by AD during certain ages [16,18], IL-16 and IL-8 levels are unlikely to be a mechanism for this effect as differences in sex by genotype were noticed in the absence not presence of APOEε4 alleles.

### Females had higher CRP levels compared to males and CRP levels were lower in APOEε4 carriers

We found that plasma and CSF levels of CRP, a widely used inflammatory and cardiovascular marker [51,52], were independently affected by sex and APOE genotype. Females, regardless of diagnosis or APOEε4 alleles, had significantly higher plasma CRP relative to males, consistent with findings in healthy individuals [53]. Higher levels of peripheral CRP may suggest higher systemic inflammation in females, which is associated with an increased risk in all-cause dementia [54]. Higher levels of serum CRP are also associated with higher levels of serum estradiol in postmenopausal healthy females [55] which suggests that sex differences in CRP levels may be partly due to sex differences in estradiol levels or other sex hormones. A recent study using the ADNI database, found that low testosterone levels was associated with higher tau pathology especially among APOEε4 carriers, regardless of sex, suggesting that testosterone maybe neuroprotective in both sexes [56]. In addition, we found that the presence of APOEε4 alleles decreased plasma and CSF CRP levels consistent with previous research in large population studies [57,58]. In our study, we also found a trend for lower levels of plasma CRP with LMCI and AD compared to CN. Recent meta-analyses did not find differences in peripheral levels of CRP in AD compared to healthy controls [59,60]. However, in participants with mild and moderate dementia only, serum CRP levels were lower compared to the cognitively healthy group [59]. In healthy individuals, higher levels of plasma CRP in midlife are associated with a higher amyloid burden later in life in males but not females [27]. However, despite this finding, higher systemic inflammation in midlife (including CRP) is associated with greater cognitive decline later in life in both sexes in healthy individuals [28]. It is important to acknowledge evidence that midlife obesity, but not later life obesity, is associated with an increased risk to develop dementia [61,62], which may be related to altered inflammation (e.g., cytokines and CRP) due to the accumulation of adipose tissue [63,64]. It is possible that sex differences in inflammation and/or obesity earlier in life have long-term effects on the transition to MCI and/or AD.

### Females had lower cytokine and immunoglobulin levels compared to males

We found some biomarkers that were affected by sex, but not diagnosis or presence of APOEε4 alleles. For example, females had lower CSF levels of ICAM1 compared to males, regardless of APOE genotype or diagnosis, but, although a trend, the opposite effect was seen in plasma. In contrast, in healthy adults (18-55 years old), serum levels of ICAM1 are lower in females compared to males [65]. ICAM1 is a type of adhesion molecule associated with microvascular endothelial activation [66] and plasma ICAM1 levels (but not CSF levels; [67]) were higher in patients with AD [67–69]. Although in the present study we did not observe a significant effect of plasma ICAM1 with diagnosis, the unadjusted P-value was 0.063 with higher levels in LMCI and AD groups. It is intriguing that females have lower CSF levels of cytokines (IL-16, IL-18), ICAM1, and immunoglobulins (IgE and IgA) but higher plasma CRP and ICAM1 levels. Although neuroinflammation is associated with AD, it may be both a product and a driver of neurodegeneration, and it may have both beneficial and detrimental roles in AD [70,71]. In AD mouse models, inflammatory cytokines (e.g., IL1β, IL-4, IL-6, IL-10, IFNγ, TNFα) can both increase amyloid beta deposition and reduce amyloid plaque pathology [72–80]. In transgenic mice, amyloid deposition is associated with low T-cell activation suggesting that the immune system is hypo-responsive to amyloid beta [81]. Thus, increases of inflammatory markers may not always be indicative of worse neuropathology or outcomes, but may be contributing to reductions in AD neuropathology. It is also possible that males and females have varying levels of beneficial vs detrimental immune responses which can differentially affect how the disease progresses between the sexes. Indeed, we found sex differences in the correlation between CSF and plasma biomarkers (CD 40, IgA, ICAM1, and IL-3), which suggests that plasma and CSF levels may be regulated differently in males and females.

### Limitations

In this exploratory study, we used two separate ADNI datasets (CSF biomarkers and plasma biomarkers) with a large overlap of individuals (85%) but different sample sizes that resulted in differences in the demographics between the datasets and power across the datasets for the analyses conducted. Because of this, the proportion of APOE or diagnosis by sex could differ across these datasets. While the proportion of sex by APOEe4 carriers did not differ substantially between the datasets, the proportion of participants in each of the diagnosis groups was not similar across datasets causing differences in statistical power to detect the interaction term of diagnosis and sex. In addition, in this cohort the proportion of participants in the different APOEε4 allele groups was correlated with diagnosis (Supplemental Table S5). Thus, a larger cohort is required to test how sex, APOE genotype, and diagnosis interact together in one model.

More generally, the ADNI cohort is not ethnically or socioeconomically diverse, being mostly composed of self-reported white (only 12 individuals were not-white) and highly educated individuals (average 15.69 years of education). As AD incidence, prevalence, and age of onset varies by ethnicity [82–84] and education [85], our conclusions may not apply to more ethnically and socially diverse populations. In addition to sex, it is possible the underlying mechanisms of AD are different depending on ethnicity. Additionally, other pathologies in these participants, such as cancer, cardiovascular disease, smoking status, or obesity may have influenced inflammatory markers and limited our interpretations.

## CONCLUSION

The current study provides evidence that sex and presence of APOEε4 alleles are associated with CSF levels of the inflammatory markers IL-16 and IL-8. We found sex differences indicating that females had lower cytokine and immunoglobulin levels but higher plasma CRP and ICAM1 levels compared to males, although the direction of the ICAM1 finding was tissue-dependent. Together, our work suggests that that presence of APOEε4 alleles can affect cytokine levels differently in males and females and the underlying pathophysiology of aging and AD may be tissue- and sex-specific.

## ACKNOWLEDGMENTS

Data collection and sharing for this project was funded by the Alzheimer’s Disease Neuroimaging Initiative (ADNI) (National Institutes of Health Grant U01 AG024904) and DOD ADNI (Department of Defense award number W81XWH-12-2-0012). ADNI is funded by the National Institute on Aging, the National Institute of Biomedical Imaging and Bioengineering, and through generous contributions from the following: AbbVie, Alzheimer’s Association; Alzheimer’s Drug Discovery Foundation; Araclon Biotech; BioClinica, Inc.; Biogen; Bristol-Myers Squibb Company; CereSpir, Inc.; Cogstate; Eisai Inc.; Elan Pharmaceuticals, Inc.; Eli Lilly and Company; EuroImmun; F. Hoffmann-La Roche Ltd and its affiliated company Genentech, Inc.; Fujirebio; GE Healthcare; IXICO Ltd.; Janssen Alzheimer Immunotherapy Research & Development, LLC.; Johnson & Johnson Pharmaceutical Research & Development LLC.; Lumosity; Lundbeck; Merck & Co., Inc.; Meso Scale Diagnostics, LLC.; NeuroRx Research; Neurotrack Technologies; Novartis Pharmaceuticals Corporation; Pfizer Inc.; Piramal Imaging; Servier; Takeda Pharmaceutical Company; and Transition Therapeutics. The Canadian Institutes of Health Research is providing funds to support ADNI clinical sites in Canada. Private sector contributions are facilitated by the Foundation for the National Institutes of Health (www.fnih.org). The grantee organization is the Northern California Institute for Research and Education, and the study is coordinated by the Alzheimer’s Therapeutic Research Institute at the University of Southern California. ADNI data are disseminated by the Laboratory for Neuro Imaging at the University of Southern California. Funding for this study was provided by a Canadian Institutes of Health Research (CHIR) grant to LAMG (PJT-148662). PDG is funded by the Alzheimer’s Association of the USA and Brain Canada with the financial support of Health Canada through the Brain Canada Research Fund (AARF-17-529705). The views expressed herein do not necessarily represent the views of the Minister of Health or the Government of Canada. AMI is funded by a University of British Columbia Four Year Doctoral Fellowship and the CIHR Frederick Banting and Charles Best Masters Research Award. We thank Elizabeth Perez for help with data organization.

## Supplemental File

**Table S1.** Linear regression results for all CSF variables investigated by sex and APOE genotype (non-carriers or carriers of APOEε4 alleles).

| *Predictors* | Cortisol Cortisol ng/ml |  |  | C Reactive Protein ug/ml |  |  | CD 40 antigen ng/ml |  |  | Interleukin 16 pg/ml |  |  | Interleukin 3 ng/ml |  |  |
| --- | --- | --- | --- | --- | --- | --- | --- | --- | --- | --- | --- | --- | --- | --- | --- |
|  | *Estimates (CI)* | *p* | *adjusted p* | *Estimates (CI)* | *p* | *adjusted p* | *Estimates (CI)* | *p* | *adjusted p* | *Estimates (CI)* | *p* | *adjusted p* | *Estimates (CI)* | *p* | *adjusted p* |
| (Intercept) | 1.43 (-7.00 – 9.87) |  |  | -3.03 (-3.84 – -2.23) |  |  | -1.10 (-1.27 – -0.93) |  |  | 0.42 (0.15 – 0.68) |  |  | -3.03 (-3.49 – -2.57) |  |  |
| AGE (years) | 0.22 (0.12 – 0.32) |  |  | 0.01 (-0.00 – 0.02) |  |  | 0.01 (0.00 – 0.01) |  |  | 0.01 (0.00 – 0.01) |  |  | 0.01 (0.00 – 0.02) |  |  |
| EDUCATION (years) | -0.21 (-0.45 – 0.03) |  |  | -0.01 (-0.03 – 0.02) |  |  | -0.00 (-0.01 – 0.00) |  |  | -0.01 (-0.01 – 0.00) |  |  | 0.00 (-0.01 – 0.01) |  |  |
| Male (ref = Female) | 1.72 (0.25 – 3.19) | 0.022 | **0.07** | -0.11 (-0.25 – 0.03) | 0.126 | 0.22 | 0.01 (-0.02 – 0.04) | 0.432 | 0.55 | 0.13 (0.07 – 0.19) |  |  | 0.08 (-0.00 – 0.16) | 0.057 | 0.15 |
| APOE4 1 or 2 alleles (ref = 0 alleles) | 0.82 (-0.55 – 2.19) | 0.241 | 0.35 | -0.22 (-0.35 – -0.09) | 0.001 | **0.009** | 0.01 (-0.02 – 0.04) | 0.59 | 0.72 | 0.08 (0.01 – 0.14) |  |  | -0.04 (-0.12 – 0.03) | 0.262 | 0.37 |
| Interaction: Male by 1 or 2 alleles |  |  |  |  |  |  |  |  |  | -0.13 (-0.22 – -0.05) | 0.003 | **0.016** |  |  |  |
| Observations | 279 |  |  | 279 |  |  | 279 |  |  | 279 |  |  | 279 |  |  |
| R<sup>2</sup> / adjusted R<sup>2</sup> | 0.095 / 0.082 |  |  | 0.055 / 0.042 |  |  | 0.125 / 0.112 |  |  | 0.122 / 0.106 |  |  | 0.076 / 0.063 |  |  |

**Table S1.** Continued
| *Predictors* | Interleukin 6 receptor ng/ml |  |  | Interleukin 8 pg/ml |  |  | Immunoglobulin A mg/ml |  |  | Intercellular Adhesion Molecule ng/ml |  |  |
| --- | --- | --- | --- | --- | --- | --- | --- | --- | --- | --- | --- | --- |
|  | *Estimates (CI)* | *p* | *adjusted p* | *Estimates (CI)* | *p* | *adjusted p* | *Estimates (CI)* | *p* | *adjusted p* | *Estimates (CI)* | *p* | *adjusted p* |
| (Intercept) | -0.27 (-0.48 – -0.06) |  |  | 1.33 (1.11 – 1.54) |  |  | -2.91 (-3.35 – -2.48) |  |  | -0.23 (-0.83 – 0.38) |  |  |
| AGE (years) | 0.00 (0.00 – 0.01) |  |  | 0.00 (0.00 – 0.01) |  |  | 0.00 (-0.00 – 0.01) |  |  | 0.01 (0.01 – 0.02) |  |  |
| EDUCATION (years) | -0.00 (-0.01 – 0.00) |  |  | 0.00 (-0.00 – 0.01) |  |  | 0.00 (-0.01 – 0.01) |  |  | -0.00 (-0.02 – 0.02) |  |  |
| Male (ref = Female) | 0.02 (-0.02 – 0.05) | 0.372 | 0.5 | 0.09 (0.04 – 0.14) | <0.001 |  | 0.21 (0.14 – 0.29) | <0.001 | **<0.001** | 0.18 (0.07 – 0.28) | 0.001 | **0.009** |
| APOE4 1 or 2 alleles (ref = 0 alleles) | 0.04 (0.00 – 0.07) | 0.025 | **0.071** | 0.03 (-0.02 – 0.09) | 0.261 |  | -0.00 (-0.07 – 0.07) | 0.898 | 0.97 | 0.08 (-0.02 – 0.18) | 0.128 | 0.22 |
| Interaction: Male by 1 or 2 alleles |  |  |  | -0.09 (-0.16 – -0.02) | 0.008 | **0.035** |  |  |  |  |  |  |
| Observations | 279 |  |  | 279 |  |  | 279 |  |  | 279 |  |  |
| R<sup>2</sup> / adjusted R<sup>2</sup> | 0.046 / 0.032 |  |  | 0.092 / 0.076 |  |  | 0.123 / 0.111 |  |  | 0.105 / 0.092 |  |  |

**Table S2.** Linear regression results for all CSF variables investigated by sex and baseline diagnosis (CN, cognitively normal; LMCI, late mild cognitive impairment; AD, Alzheimer’s disease). There were no significant interactions between diagnosis and sex.

| Predictors | Cortisol Cortisol ng/ml |  |  | C Reactive Protein ug/ml |  |  | CD 40 antigen ng/ml |  |  | Interleukin 16 pg/ml |  |  | Interleukin 3 ng/ml |  |  |
| --- | --- | --- | --- | --- | --- | --- | --- | --- | --- | --- | --- | --- | --- | --- | --- |
|  | Estimates (CI) | p | adjusted p | Estimates (CI) | p | adjusted p | Estimates (CI) | p | adjusted p | Estimates (CI) | p | adjusted p | Estimates (CI) | p | adjusted p |
| (Intercept) | 1.49 (-6.96 – 9.95) |  |  | -3.12 (-3.94 – -2.30) |  |  | -1.08 (-1.25 – -0.91) |  |  | 0.51 (0.26 – 0.77) |  |  | -3.03 (-3.49 – -2.57) |  |  |
| AGE (years) | 0.22 (0.12 – 0.32) |  |  | 0.01 (-0.00 – 0.02) |  |  | 0.01 (0.00 – 0.01) |  |  | 0.01 (0.00 – 0.01) |  |  | 0.01 (0.00 – 0.02) |  |  |
| EDUCATION (years) | -0.23 (-0.47 – 0.01) |  |  | -0.00 (-0.03 – 0.02) |  |  | -0.00 (-0.01 – 0.00) |  |  | -0.01 (-0.01 – 0.00) |  |  | 0.00 (-0.01 – 0.01) |  |  |
| Male (ref = Female) | 1.55 (0.07 – 3.04) | 0.041 | 0.18 | -0.10 (-0.24 – 0.05) | 0.19 | 0.42 | 0.01 (-0.02 – 0.04) | 0.51 | 0.65 | 0.06 (0.02 – 0.11) | 0.007 | **0.055** | 0.08 (-0.00 – 0.16) | 0.052 | 0.18 |
| Diagnosis (ref = CN) |  | 0.24 | 0.49 |  | 0.15 | 0.36 |  | 0.067 | 0.2 |  | 0.4 | 0.61 |  | 0.5 | 0.65 |
| LMCI | 1.23 (-0.43 – 2.88) |  |  | -0.15 (-0.31 – 0.01) |  |  | 0.01 (-0.03 – 0.04) |  |  | 0.01 (-0.05 – 0.06) |  |  | -0.04 (-0.13 – 0.05) |  |  |
| AD | 0.10 (-1.83 – 2.02) |  |  | -0.15 (-0.34 – 0.03) |  |  | -0.03 (-0.07 – 0.00) |  |  | -0.03 (-0.09 – 0.03) |  |  | -0.06 (-0.17 – 0.04) |  |  |
| Observations | 279 |  |  | 279 |  |  | 279 |  |  | 279 |  |  | 279 |  |  |
| R<sup>2</sup> / adjusted R<sup>2</sup> | 0.100 / 0.084 |  |  | 0.030 / 0.012 |  |  | 0.141 / 0.125 |  |  | 0.099 / 0.082 |  |  | 0.077 / 0.060 |  |  |

**Table S2.** Continued
| Predictors | Interleukin 6 receptor ng/ml |  |  | Interleukin 8 pg/ml |  |  | Immunoglobulin A mg/ml |  |  | Intercellular Adhesion Molecule ng/ml |  |  |
| --- | --- | --- | --- | --- | --- | --- | --- | --- | --- | --- | --- | --- |
|  | Estimates (CI) | p | adjusted p | Estimates (CI) | p | adjusted p | Estimates (CI) | p | adjusted p | Estimates (CI) | p | adjusted p |
| (Intercept) | -0.22 (-0.43 – -0.01) |  |  | 1.35 (1.14 – 1.56) |  |  | -2.92 (-3.35 – -2.48) |  |  | -0.19 (-0.80 – 0.42) |  |  |
| AGE (years) | 0.00 (0.00 – 0.01) |  |  | 0.00 (0.00 – 0.01) |  |  | 0.00 (-0.00 – 0.01) |  |  | 0.01 (0.01 – 0.02) |  |  |
| EDUCATION (years) | -0.00 (-0.01 – 0.00) |  |  | 0.00 (-0.00 – 0.01) |  |  | 0.00 (-0.01 – 0.01) |  |  | -0.00 (-0.02 – 0.02) |  |  |
| Male (ref = Female) | 0.02 (-0.02 – 0.05) | 0.42 | 0.61 | 0.04 (0.01 – 0.08) | 0.019 | 0.12 | 0.21 (0.14 – 0.29) | <0.001 | **<0.001** | 0.17 (0.06 – 0.28) | 0.002 | **0.026** |
| Diagnosis (ref = CN) |  | 0.46 | 0.65 |  | 0.85 | 0.96 |  | 0.99 | 0.99 |  | 0.54 | 0.67 |
| LMCI | 0.01 (-0.04 – 0.05) |  |  | 0.01 (-0.03 – 0.05) |  |  | 0.00 (-0.08 – 0.09) |  |  | 0.06 (-0.06 – 0.18) |  |  |
| AD | -0.02 (-0.07 – 0.03) |  |  | 0.01 (-0.03 – 0.06) |  |  | -0.01 (-0.11 – 0.09) |  |  | 0.02 (-0.12 – 0.16) |  |  |
| Observations | 279 |  |  | 279 |  |  | 279 |  |  | 279 |  |  |
| R<sup>2</sup> / adjusted R<sup>2</sup> | 0.034 / 0.016 |  |  | 0.062 / 0.045 |  |  | 0.124 / 0.107 |  |  | 0.101 / 0.085 |  |  |

**Table S3.** Linear regression results for all plasma variables investigated by sex and APOE genotype (non-carriers or carriers of APOEε4 alleles).

| Predictors | Cortisol Cortisol ng/ml |  |  | C Reactive Protein ug/ml |  |  | CD 40 antigen ng/ml |  |  | Interleukin 16 pg/ml |  |  | Interleukin 3 ng/ml |  |  |
| --- | --- | --- | --- | --- | --- | --- | --- | --- | --- | --- | --- | --- | --- | --- | --- |
|  | Estimates (CI) | p | adjusted p | Estimates (CI) | p | adjusted p | Estimates (CI) | p | adjusted p | Estimates (CI) | p | adjusted p | Estimates (CI) | p | adjusted p |
| (Intercept) | 2.06 (1.92 – 2.19) |  |  | 0.46 (-0.05 – 0.97) |  |  | -0.52 (-0.65 – -0.39) |  |  | 2.42 (2.27 – 2.58) |  |  | -1.86 (-2.16 – -1.57) |  |  |
| AGE (years) | 0.00 (-0.00 – 0.00) |  |  | 0.00 (-0.00 – 0.01) |  |  | 0.01 (0.00 – 0.01) |  |  | 0.00 (0.00 – 0.00) |  |  | 0.00 (-0.00 – 0.00) |  |  |
| EDUCATION (years) | -0.00 (-0.00 – 0.00) |  |  | -0.03 (-0.04 – -0.01) |  |  | -0.00 (-0.01 – 0.00) |  |  | -0.01 (-0.01 – -0.00) |  |  | 0.01 (0.00 – 0.02) |  |  |
| Male (ref = Female) | 0.01 (-0.01 – 0.04) | 0.26 | 0.57 | -0.13 (-0.22 – -0.04) | 0.006 | **0.048** | -0.04 (-0.07 – -0.01) |  |  | 0.01 (-0.02 – 0.04) | 0.50 | 0.79 | -0.02 (-0.07 – 0.04) | 0.57 | 0.8 |
| APOE4 1 or 2 alleles (ref = 0 alleles) | 0.02 (-0.00 – 0.04) | 0.11 | 0.29 | -0.31 (-0.39 – -0.22) | <0.001 | **<0.001** | -0.03 (-0.07 – 0.00) |  |  | -0.01 (-0.04 – 0.01) | 0.31 | 0.66 | 0.02 (-0.03 – 0.07) | 0.43 | 0.67 |
| Interaction: Male by 1 or 2 alleles |  |  |  |  |  |  | 0.05 (0.01 – 0.10) | **0.022** | 0.11 |  |  |  |  |  |  |
| Observations | 527 |  |  | 526 |  |  | 527 |  |  | 527 |  |  | 527 |  |  |
| R<sup>2</sup> / adjusted R<sup>2</sup> | 0.013 / 0.005 |  |  | 0.123 / 0.117 |  |  | 0.126 / 0.118 |  |  | 0.042 / 0.033 |  |  | 0.011 / 0.004 |  |  |

**Table S3.** Continued
| Predictors | Interleukin 6 receptor ng/ml |  |  | Interleukin 8 pg/ml |  |  | Immunoglobulin A mg/ml |  |  | Intercellular Adhesion Molecule ng/ml |  |  | Interleukin 18 pg/ml |  |  | Immunoglobulin E ng/ml |  |  |
| --- | --- | --- | --- | --- | --- | --- | --- | --- | --- | --- | --- | --- | --- | --- | --- | --- | --- | --- |
|  | Estimates (CI) | p | adjusted p | Estimates (CI) | p | adjusted p | Estimates (CI) | p | adjusted p | Estimates (CI) | p | adjusted p | Estimates (CI) | p | adjusted p | Estimates (CI) | p | adjusted p |
| (Intercept) | 1.43 (1.29 – 1.57) |  |  | 0.79 (0.60 – 0.99) |  |  | 0.48 (0.25 – 0.71) |  |  | 1.89 (1.75 – 2.04) |  |  | 2.42 (2.24 – 2.59) |  |  | 1.42 (0.88 – 1.97) |  |  |
| AGE (years) | 0.00 (-0.00 – 0.00) |  |  | 0.00 (0.00 – 0.01) |  |  | 0.00 (-0.00 – 0.00) |  |  | 0.00 (0.00 – 0.00) |  |  | -0.00 (-0.00 – 0.00) |  |  | 0.00 (-0.00 – 0.01) |  |  |
| EDUCATION (years) | -0.00 (-0.01 – 0.00) |  |  | 0.00 (-0.00 – 0.01) |  |  | -0.00 (-0.01 – 0.01) |  |  | 0.00 (-0.00 – 0.00) |  |  | -0.00 (-0.01 – 0.00) |  |  | 0.01 (-0.01 – 0.02) |  |  |
| Male (ref = Female) | -0.02 (-0.05 – -0.00) | 0.049 | 0.14 | -0.05 (-0.10 – 0.00) |  |  | 0.02 (-0.02 – 0.06) | 0.371 | 0.67 | -0.04 (-0.06 – -0.01) | 0.008 | **0.051** | 0.06 (0.03 – 0.09) | <0.001 | **0.001** | 0.24 (0.14 – 0.33) | <0.001 | **<0.001** |
| APOE4 1 or 2 alleles (ref = 0 alleles) | 0.00 (-0.02 – 0.03) | 0.70 | 0.84 | -0.08 (-0.13 – -0.02) |  |  | -0.02 (-0.06 – 0.02) | 0.387 | 0.67 | 0.01 (-0.02 – 0.03) | 0.586 | 0.80 | -0.04 (-0.07 – -0.01) | 0.019 | 0.10 | 0.01 (-0.09 – 0.10) | 0.91 | 0.94 |
| Interaction: Male by 1 or 2 alleles |  |  |  | 0.07 (0.00 – 0.14) | 0.046 | 0.14 |  |  |  |  |  |  |  |  |  |  |  |  |
| Observations | 527 |  |  | 527 |  |  | 527 |  |  | 527 |  |  | 527 |  |  | 527 |  |  |
| R<sup>2</sup> / adjusted R<sup>2</sup> | 0.011 / 0.004 |  |  | 0.036 / 0.026 |  |  | 0.019 / 0.012 |  |  | 0.008 / 0.000 |  |  | 0.038 / 0.031 |  |  | 0.052 / 0.045 |  |  |

**Table S4.** Linear regression results for all plasma variables investigated by sex and baseline diagnosis (CN, cognitively normal; LMCI, late mild cognitive impairment; AD, Alzheimer’s disease). There were no significant interactions between diagnosis and sex.

|  | Cortisol ng/ml |  |  | C Reactive Protein ug/ml |  |  | CD 40 antigen ng/ml |  |  | Interleukin 16 pg/ml |  |  | Interleukin 3 ng/ml |  |  |
| --- | --- | --- | --- | --- | --- | --- | --- | --- | --- | --- | --- | --- | --- | --- | --- |
| Predictors | Estimates (CI) | p | adjusted p | Estimates (CI) | p | adjusted p | Estimates (CI) | p | adjusted p | Estimates (CI) | p | adjusted p | Estimates (CI) | p | adjusted p |
| (Intercept) | 2.09 (1.95 – 2.22) |  |  | 0.33 (-0.21 – 0.88) |  |  | -0.55 (-0.68 – -0.42) |  |  | 2.48 (2.31 – 2.65) |  |  | -1.78 (-2.08 – -1.47) |  |  |
| AGE (years) | 0.00 (-0.00 – 0.00) |  |  | 0.01 (0.00 – 0.01) |  |  | 0.01 (0.00 – 0.01) |  |  | 0.00 (0.00 – 0.00) |  |  | 0.00 (-0.00 – 0.00) |  |  |
| EDUCATION (years) | -0.00 (-0.00 – 0.00) |  |  | -0.02 (-0.04 – -0.01) |  |  | -0.00 (-0.00 – 0.00) |  |  | -0.01 (-0.01 – -0.00) |  |  | 0.01 (0.00 – 0.02) |  |  |
| Male (ref = Female) | 0.02 (-0.01 – 0.04) | 0.18 | 0.44 | -0.12 (-0.21 – -0.03) | 0.012 | **0.056** | -0.01 (-0.03 – 0.01) | 0.38 | 0.59 | 0.01 (-0.02 – 0.04) | 0.39 | 0.57 | -0.02 (-0.07 – 0.04) | 0.56 | 0.75 |
| Diagnosis (ref = CN) |  | 0.0009 | **0.010** |  | 0.007 | **0.056** |  | 0.016 | **0.067** |  | 0.009 | **0.054** |  | 0.044 | 0.17 |
| LMCI | -0.02 (-0.07 – 0.02) |  |  | -0.27 (-0.44 – -0.10) |  |  | -0.02 (-0.06 – 0.02) |  |  | -0.08 (-0.13 – -0.03) |  |  | -0.04 (-0.13 – 0.06) |  |  |
| AD | 0.03 (-0.02 – 0.08) |  |  | -0.24 (-0.43 – -0.05) |  |  | 0.02 (-0.02 – 0.07) |  |  | -0.08 (-0.14 – -0.02) |  |  | -0.11 (-0.21 – -0.00) |  |  |
| Observations | 527 |  |  | 526 |  |  | 527 |  |  | 526 |  |  | 527 |  |  |
| R<sup>2</sup> / adjusted R<sup>2</sup> | 0.034 / 0.024 |  |  | 0.061 / 0.052 |  |  | 0.131 / 0.123 |  |  | 0.048 / 0.039 |  |  | 0.022 / 0.013 |  |  |

**Table S4.** Continued
|  | Interleukin 6 receptor ng/ml |  |  | Interleukin 8 pg/ml |  |  | Immunoglobulin A mg/ml |  |  | Intercellular Adhesion Molecule ng/ml |  |  | Interleukin 18 pg/ml |  |  | Immunoglobulin E ng/ml |  |  |
| --- | --- | --- | --- | --- | --- | --- | --- | --- | --- | --- | --- | --- | --- | --- | --- | --- | --- | --- |
| Predictors | Estimates (CI) | p | adjusted p | Estimates (CI) | p | adjusted p | Estimates (CI) | p | adjusted p | Estimates (CI) | p | adjusted p | Estimates (CI) | p | adjusted p | Estimates (CI) | p | adjusted p |
| (Intercept) | 1.47 (1.33 – 1.61) |  |  | 0.73 (0.54 – 0.93) |  |  | 0.43 (0.20 – 0.67) |  |  | 1.90 (1.75 – 2.05) |  |  | 2.33 (2.15 – 2.51) |  |  | 1.44 (0.89 – 2.00) |  |  |
| AGE (years) | 0.00 (-0.00 – 0.00) |  |  | 0.00 (0.00 – 0.01) |  |  | 0.00 (-0.00 – 0.00) |  |  | 0.00 (-0.00 – 0.00) |  |  | 0.00 (-0.00 – 0.00) |  |  | 0.00 (-0.00 – 0.01) |  |  |
| EDUCATION (years) | -0.00 (-0.01 – 0.00) |  |  | 0.00 (-0.00 – 0.01) |  |  | -0.00 (-0.01 – 0.01) |  |  | 0.00 (-0.00 – 0.01) |  |  | -0.00 (-0.01 – 0.00) |  |  | 0.01 (-0.01 – 0.02) |  |  |
| Male (ref = Female) | -0.02 (-0.05 – 0.00) | 0.063 | 0.19 | -0.01 (-0.04 – 0.03) | 0.65 | 0.79 | 0.02 (-0.02 – 0.06) | 0.42 | 0.59 | -0.03 (-0.06 – -0.01) | 0.011 | **0.056** | 0.06 (0.03 – 0.09) | <0.001 | **0.004** | 0.24 (0.14 – 0.33) | <0.001 | **<0.001** |
| Diagnosis (ref = CN) |  | 0.34 | 0.56 |  | 0.11 | 0.30 |  | 0.72 | 0.82 |  | 0.063 | 0.19 |  | 0.32 | 0.56 |  | 0.88 | 0.94 |
| LMCI | -0.03 (-0.08 – 0.01) |  |  | -0.04 (-0.10 – 0.02) |  |  | 0.03 (-0.05 – 0.10) |  |  | -0.01 (-0.06 – 0.03) |  |  | 0.04 (-0.01 – 0.10) |  |  | -0.00 (-0.18 – 0.17) |  |  |
| AD | -0.03 (-0.08 – 0.02) |  |  | -0.00 (-0.07 – 0.07) |  |  | 0.02 (-0.07 – 0.10) |  |  | 0.02 (-0.03 – 0.08) |  |  | 0.03 (-0.03 – 0.10) |  |  | -0.03 (-0.23 – 0.16) |  |  |
| Observations | 527 |  |  | 527 |  |  | 527 |  |  | 527 |  |  | 527 |  |  | 527 |  |  |
| R<sup>2</sup> / adjusted R<sup>2</sup> | 0.015 / 0.006 |  |  | 0.029 / 0.020 |  |  | 0.008 / -0.002 |  |  | 0.029 / 0.019 |  |  | 0.032 / 0.023 |  |  | 0.053 / 0.043 |  |  |

**Table S5.**
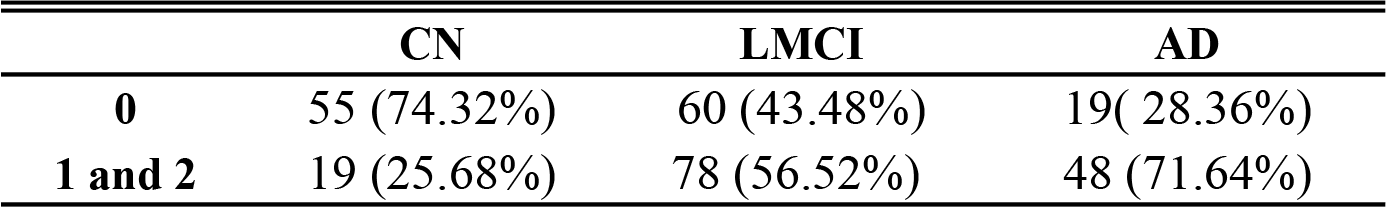
Contingency table for the distribution of the APOEε4 alleles (0 or 1 and 2 ε4 alleles) in each of the diagnosis groups (CN, cognitively normal; LMCI, late mild cognitive impairment; AD, Alzheimer’s disease).

|  | CN | LMCI | AD |
| --- | --- | --- | --- |
| **0** | 55 (74.32%) | 60 (43.48%) | 19( 28.36%) |
| **1 and 2** | 19 (25.68%) | 78 (56.52%) | 48 (71.64%) |

